# Toxicity, transfer and depuration of anatoxin-a (cyanobacterial neurotoxin) in medaka fish exposed by single-dose gavage

**DOI:** 10.1101/868737

**Authors:** Simon Colas, Charlotte Duval, Benjamin Marie

## Abstract

The proliferations of cyanobacteria are increasingly prevalent in warm and nutrient-enriched waters and occur in many rivers and water bodies due especially to eutrophication. The aim of this work is to study in female medaka fish the toxicity, the transfer and the depuration of the anatoxin-a, a neurotoxin produced by benthic cyanobacterial biofilms. This work will provide answers regarding acute toxicity induced by single gavage by anatoxin-a and to the risks of exposure by ingestion of contaminated fish flesh, considering that data on these aspects remain particularly limited.

The oral LD_50_ of a single dose of (±)-anatoxin-a was determined at 11.50 µg.g^−1^. First of all, lethal dose (100% from 20 µg.g^−1^) provokes rapid respiratory paralysis (in 1-2 min) of the fish inducing the death by asphyxia. Noticeably, no death nor apparent neurotoxicologic effect occurred during the experimentation period for the 45 fish exposed to a single sub-acute dose of (±)-anatoxin-a corresponding to the no observable adverse effect level (NOAEL = 6.67 µg.g^−1^). Subsequently, the toxico-kinetics of the (±)-anatoxin-a was observed in the guts, the livers and the muscles of female medaka fish for 10 days.

In parallel, a protocol for extraction of anatoxin-a has been optimized beforehand by testing 3 different solvents on several matrices, the extraction with 75% methanol + 0.1% formic acid appearing to be the most efficient. Anatoxin-a was quantified by high-resolution qTOF mass spectrometry coupled upstream to a UHPLC chromatographic chain. The toxin could not be detected in the liver after 12 h, and in the gut and muscle after 3 days. The mean clearance rates of (±)-anatoxin-a calculated after 12 h are above 58%, 100% and 90% for the guts, the livers and the muscles, respectively. Non-targeted metabolomics investigations performed on the fish liver indicates that the single sub-acute exposure by gavage induces noticeable metabolome dysregulations, including important phospholipid decreases, with an organism recovery period of above 12-24h. Overall, the medaka fish do not appear to accumulate (±)-anatoxin-a and to largely recover after 24h following a single sub-acute oral liquid exposure at the NOAEL.

## Introduction

Worldwide proliferation of cyanobacterial blooms constitutes a serious environmental and economic problem that menaces wildlife and human health. Moreover, many cyanobacteria are able to produce potent hepatotoxins such as microcystin, cylindrospermopsin and nodularin, and/or neurotoxins such as anatoxins-a, homo-anatoxin-a, anatoxin-a(s) and saxitoxin (Sivonen & Jones, 1999). Harmful cyanobacterial blooms in lakes have been described for decades; in rivers, however, first reports of animal deaths from toxic benthic cyanobacteria in many regions occurred only in the last 20 years (Wood et al., 2007). Since toxic benthic cyanobacteria in rivers have been documented in many countries and benthic taxa have been found to produce a large spectrum of those cyanotoxins, though most animal fatalities reported in relation to benthic cyanobacteria were associated with the presence of anatoxin-a (ANTX) and/or homoanatoxin-a (HANTX) (for review Quiblier et al., 2013).

Anatoxin-a and homoanatoxin-a are potent neurotoxins produced by some planktonic and benthic strains of the genera *Anabaena, Oscillatoria, Aphanizomenon* and *Cylindrospermum* (Bouma-Gerson et al., 2018). Indeed, numerous fatal intoxications of animals, by anatoxin-a and homoanatoxin-a, have been reported all over the world (Gugger et al., 2005; Puschner et al., 2010; Wood et al., 2007). These alkaloids bind tightly to the nicotinic acetylcholine receptor, in the sub-nanomolar range, and thus provoke the death of animals almost immediately after ingestion (Wonnacott and Gallagher, 2006). Anatoxin-a, as a competitive agonist of acetylcholine can bind it specific membrane receptors, notably at neuromuscular junctions, affecting signal transmission between neurons and muscles, causing muscle cell overstimulation (Carmichael, 1994; Aráoz et al. 2010). Acute effects in vertebrates include a rapid loss of coordination, a decreased of locomotor activity, a paralysis of the peripheral skeletal and respiratory muscles, causing symptoms such as loss of coordination, twitching, irregular breathing, tremors, altered gait and convulsions before death by acute asphyxia induced by respiratory arrest (Dittmann and Wiegand, 2006).

It is maybe because anatoxin-a is very unstable and labile in the water (Stevens and Krieger, 1991) and because no chronic effects have been described in mammals so far (Fawell et al., 1999), that this toxin has been considered of less environmental concern comparing to other cyanotoxins. Despite the acute neurotoxic effects of anatoxin-a, the consequence of the proliferation of anatoxins-a producing cyanobacterial biofilms on ecosystem health and aquatic organisms remains largely unknown. Despite, toxic benthic cyanobacterial mats have been associated with decreased macro-invertebrate diversity (Aboal et al., 2002), few studies have investigated the genuine toxicological effects of these compounds on aquatic organisms (Carneiro et al., 2015; Osswald et al., 2007b; Anderson et al., 2018). In some experiments performed on fishes, such as carps and goldfish, behaviourial defects such as rapid opercular movement, abnormal swimming (Osswald et al., 2007a) and muscle rigidity (Carmichael et al., 1975) were observed. Oberemm et al. (1999) described also the alterations in heart rates in zebrafish embryos after an exposure to anatoxin-a. Thus a scarcity of information regarding its capability of aquatic species to bio-concentrate and bioaccumulation anatoxin-a when administrated by natural oral pathways, and its subsequent potential toxicological impacts on organisms still exists. In a previous study, Osswald and co-workers (2007b) found that anatoxin-a may be bioaccumulated by carps in significant levels (0.768 μg.g^−1^ of carp weight). Whether this may have an impact in aquatic food webs is not yet known.

In this study, we wanted to study the toxico-kinetics of the anatoxin-a and its consecutive possibility of accumulation of in fish tissues. We use medaka model fish in order to be able to administrate, under controlled conditions by a single gavage, predetermined dose of anatoxin-a in order to determine the dose-response toxicity parameters and to follow the assimilation/depuration processes of fish gavaged with a no observable adverse effect level (NOAEL) dose during the 10 following days. As there is still a lack of reference and standardized protocol for anatoxins-a extraction from biological matrix, such as fish tissues, and for quantification analysis, we have also tested three different extraction procedures inspired by previously published works (Triantis et al., 2016) and describe a high accuracy detection method developed on UHPLC coupled HR-qTOF mass spectrometer using TASQ software. This present work provides significant outcomes for the investigation of fish contamination by anatoxin-a.

## Material and methods

### Chemicals

In solution certified (+)-Anatoxin-a and dry (±)-Anatoxin-a were purchased from CRM-NRC (Canada) and Abcam (UK), respectively. The purity and the concentration of the daily reconstituted (±)-Anatoxin-a in ultra-pure water was initially checked by LC–MS/MS as described in the following protocol. UHPLC-MS grade methanol and acetonitrile were purchased from Bio TechnoFix (France). Proteomics grade formic acid was purchased from Sigma-Aldrich (Germany).

### Cyanobacteria cultures and Anatoxin-a extraction procedure test

Three monoclonal non-axenic cultures of *Phomidium* sp. (PMC 1001.17, 1007.17 and 1008.17) maintained at 25°C in 15-mL vessels with Z8 media in the PMC (Paris Museum Collection) of living cyanobacteria. Larger volume of all strains was simultaneously cultivated during one month in triplicates in 250 mL Erlenmeyer vessels at 25°C using a Z8 medium with a 16h:8h light/dark cycle (60 µmol.m^−2^.s^−1^). Cyanobacterial cells were centrifuged (at 4,000 g for 10 min), then freeze-dried and stored at −80°C prior to anatoxin-a extraction. The lyophilized cells were weighted, sonicated 2 min in a constant ratio of 100 µL of solvent for 1 mg of dried biomass, centrifuged at 4°C (12,000 g; 10 min), then the supernatant was collected and directly analysed by mass spectrometry. We have presently tested in triplicates the extraction efficiency of three different solvent mixtures already propositioned for anatoxin-a extraction (Bogialli et al., 2006; Rellan et al., 2007; Triantis et al., 2016; Haddad et al., 2019), comprising: a pure water solution acidified with 0.1% formic acid (“Water” extraction), a 25-% acetonitrile solution acidified with 0.1% formic acid (“Acetonitrile” extraction), and a 75-% methanol solution acidified with 0.1% formic acid (“Methanol” extraction). The efficiency of the extraction procedure was significantly determined according to Dunn’s post-hoc test performed after Krustal-Wallis non-parametric tests process on R software.

### Fish experimentation

Experiments on medaka fish was conducted according to European community good practices, validated by the ethical “comité Cuvier” (Author. APAFiS#19417-2019022711043436 v4) and under the supervision of accredited personnel (B.M.). Adult female medaka fishes (*Oryzias latipes*) of the inbred cab strain and of above 1 ± 0.1 g of wet weight were used in all experiments. They were raised in 20-L glass aquaria filled with a continuously aerated mixture of tap water and reverse osmosis filtered water (1/3–2/3, respectively), which was changed once a week. Fish were maintained at 25 ± 1 °C, with a 12 h:12 h light:dark standard cycle.

Fish were individually anesthetized in 0.1% tricaine methane sulfonate (MS-222; Sigma, St. Louis, MO), and then, briefly, 2 μL of a (±)-anatoxin-a mixture containing from 0.2 to 20 μg of (±)-anatoxin-a in water saturated with phenol red dyes was administrated by gavage performed with a smooth plastic needle. Control fishes were gavaged with 2 μL of water saturated with phenol red dyes. For all individual, the efficiency of the gavage uptakes was carefully checked according to the total lack of phenol red release from the mouth or the gill opercula of the fish, otherwise the individual was immediately sacrificed. The fish was then instantaneously placed in fresh water tanks and individually observed during 30 min in order to detect any behavioural sign of anatoxin-a neurotoxicity, that comprises: paralysis, decrease of breathing activity through opercular movements, locomotors activity or buoyancy default. Indeed, when gavaged with none toxic dose or control mixture, fish promptly recovers from anaesthesia within a minute and rapidly normal swimming activity (in less than 3-5 minutes). But, alternatively, the toxicological effects of toxic doses of (±)-anatoxin-a induce immediate neuro-muscular pathology and provoke a full breathing stop. After 30 min of observation, the individual was declared as “dead” as no recover was observed and the experiment was then concluded, then all fishes were anesthetized in 0.1% tricaine methane sulfonate (MS-222) and euthanized. The medium lethal dose (LD_50_) and the no observable adverse effect limit (NOAEL) were calculated from toxicological results by logistic regression after log transformation of the concentration values using R software.

For toxico-kinetics investigations, adult female medaka fish were similarly gavaged individually by a single NOAEL dose, then placed in fresh water and collected after 1h, 3h, 6h, 12h, 24h, 3d, 7d or 10d of maintaining under classical conditions. Accordingly, all fishes were anesthetized in 0.1% tricaine methane sulfonate, sacrificed, dissected, and the whole gut, the liver and the muscles were sampled and flash-frozen in liquid nitrogen, and kept frozen at - 80°C prior to analysis.

### Anatoxin-a and metabolite extraction from fish tissues

The fish tissues were weighted then sonicated 2 min in a constant ratio of 10 µL of 75-% methanol solution acidified with 0.1% formic acid for 1 mg of wet biomass for guts and livers, and of lyophilised biomass for muscles, grinded into a fine powder on Tissue-lyser (with 5 mm steel beads, Qiagen), centrifuged at 4°C (12,000 g; 10 min); then the supernatant was collected and directly analysed by mass spectrometry. The efficiency of the extraction was estimated according to the recovery rate determined in triplicats by injecting known amount of (±)-anatoxin-a to negative samples before extraction (method A) or just before the mass spectrometry analysis (method B).

### Anatoxin-a detection and quantification

Ultra high performance liquid chromatography (UHPLC) was performed on 2 µL of each of the metabolite extracts using a Polar Advances II 2.5 pore C_18_ column (Thermo) at a 300 µL.min^−1^ flow rate with a linear gradient of acetonitrile in 0.1% formic acid (5 to 90 % in 21 min). The metabolite contents were analyzed in triplicate for each strain using an electrospray ionization hybrid quadrupole time-of-flight (ESI-QqTOF) high resolution mass spectrometer (Maxis II ETD, Bruker) at 2 Hz speed on simple MS mode and subsequently on broad-band Collision Ion Dissociation (bbCID) MS/MS mode on the 50–1500 m/z rang. Calibrants, composed by serial dilutions of (+)-anatoxin-a, and negative control (phenylalanine) were analysed similarly. The raw data were automatically process with the TASQ 1.4 software for internal recalibration (< 0.5 ppm for each sample, as an internal calibrant of Na formate was injected at the beginning of each analysis) for global screening and quantification, and molecular featuring, respectively. Then, the automatic screening and quantification of anatoxin-a was performed with threshold parameters set to recommended default value for the Maxis II mass spectrometer (*Δ* RT < 0.4 s, *Δ m/z* < 3 ppm, mSigma < 50 and S/N < 5). Quantification was performed according to the integration of the area under the peaks and calibration curve was performed with certified standards.

### Analysis of liver metabolomes

Metabolites composition of the fish livers were analysed by injection of 2 μL of the 75% methanol extracts on an UHPLC (ELUTE, Bruker) coupled with a high-resolution mass spectrometer (ESI-Qq-TOF Compact, Bruker) at 2 Hz speed on simple MS mode and subsequently on broad-band Collision Ion Dissociation (bbCID) or autoMS/MS mode on the 50–1,500 *m/z* rang. The analyte annotations were performed according to precise mass and isotopic and fragmentation MS/MS patterns, as previously described (Kim Tiam et al., 2019). The feature peak list was generated from recalibrated MS spectra (< 0.5 ppm for each sample, as an internal calibrant of Na formate was injected at the beginning of each analysis) within a 1-15 min window of the LC gradient, with a filtering of 5,000 count of minimal intensity, a minimal occurrence in at least 50% of all samples, and combining all charge states and related isotopic forms using MetaboScape 4.0 software (Bruker). The intensity data table of the 591 extracted analytes was further treated using MetaboAnalyst 4 tool (Chong et al., 2019) for Pareto’s normalization, ANOVA, PCA and PLS-DA, and data representation by heatmap with hierarchical clustering, loading plots and box plots.

Unsupervised PCA models were first used to evaluate the divide between experimental groups, while supervised PLS-DA models allowed us to increase the separation between sample classes and to extract information on discriminating metabolites. The PLS-DA allowed the determination of discriminating metabolites using the analytes score values of the variable importance on projection (VIP) indicating the respective contribution of a variable to the discrimination between all of the experimental classes of samples. The higher score being in agreement with a strongest discriminatory ability and thus constitutes a criterion for the selection of the analytes as discriminative components. The PLS models were tested for over fitting with methods of permutation tests. The descriptive, predictive and consistency performance of the models was determined by R^2^, Q^2^ values and permutation test results (n = 100), respectively.

## Results

### Anatoxin-a extraction and analysis

Anatoxin-a specific mass spectrometry detection and quantification was automatically determined with high accuracy using TASQ (Bruker, Germany) from raw data generated by LC-MS/MS system according to the observation of analytes signal exhibiting targeted precise molecular mass, retention time, isotopic pattern and fragmentation ions (Fig. 1A-B). This approach allows to discriminate both (+)-anatoxin-a and (-)-anatoxin-a isomers according to their respective retention times (Fig. 1C), as well as phenylalanine (Fig. 1D) used as negative control, that does not exhibit any signal interaction with the anatoxin-a quantification. Standard solutions containing certified quantity of (+)-anatoxin-a was diluted in ultra-pure water in the range of 5 μg.mL^−1^ - 2 ng.mL^−1^ and used for calibration with good linearity of the calibration curve exhibiting a correlation coefficient with R^2^ = 0.99326 (Fig. 1E). Quantification of the (+)-anatoxin-a was performed according to the area-under-the-curve signal that was automatically integrated and process by the software that provide calculation details comprising all diagnostic elements in the report table generate for each analysis (Fig 1F).

**Figure 1.**
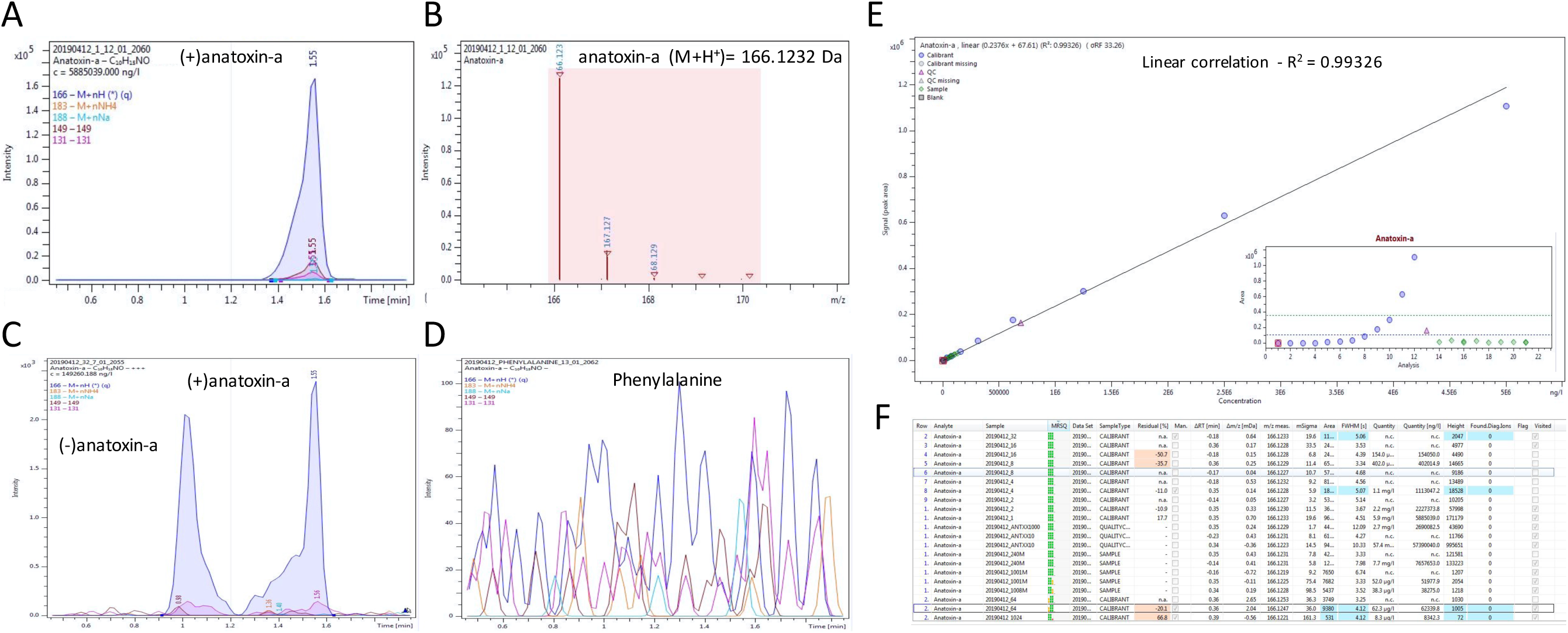
Anatoxin-a detection and quantification on HR qTOF mass spectrometer coupled to an UHPLC. The screening of (+)-anatoxin-a was performed according to the specific detection of analytes presenting accurate mass of the parent ion (M+H^+^= 166.1232 ± 0.001 Da), accurate retention time (1.55 ± 0.2 min, accurate isotopic pattern (mSigma < 50) and co-detection of characteristic fragment ions (149.096 ± 0.001 and 131.086 ± 0.001 Da) (A-B). The similar analyses of the (±)-anatoxin-a can discriminate the (-) and the (+)-anatoxin-a that exhibits distinct retention times (1.1 and 1.6 min, respectively) (C). No signal was detected with phenylalanine (D). Calibration curve performed with 12 serial dilutions of certified standard of (+)-anatoxin-a (NRC, Canada) presenting applicable correlation factor (E). Example of screening diagnostic table provided by TASQ® software for (+)-anatoxin-a quantification summarizing all qualitative and quantitative parameters for each sample (F).

Anatoxin-a extraction efficiency was assayed in triplicates with 3 different *Phormidium* strains using rather acetonitrile, methanol of water solvent solutions. In our hands, the 75% methanol acidified with 0.1% formic acid present significantly higher efficiency for anatoxin-a extraction (Table 1) and was further performed for the fish tissue extraction. Instrument and method detection and quantitation limits (LOD and LOQ) and recovery rate were determined by using triplicate injections of the control different fish tissues spiked with of determined doses of (+)-anatoxin-a that were administrated before the extraction (method A), or just before the sample analysis by mass spectrometry (method B) (table 2). The LOD and LOQ of the technics was estimated to be in the same rang as those previously described (Triantis et al., 2016). These investigations also show that anatoxin-a extraction and detection recovery rate vary from 25-78% and 51-126%, respectively, according to the tissue analysis with the guts and the muscles presenting the best and the worth recovery scores, respectively.

**Table 1.** Efficiency of different protocols for anatoxin-a extraction from different cyanobacterial strains. ^a-b-c^: indicate group results in a decreasing order of Dunn’s post-hoc test performed after Krustal-Wallis non-parametric tests.

| *Phormidium* strain | Extraction solvent |  |  |
| --- | --- | --- | --- |
|  | Acetone | Water | Methanol |
| PMC 1001.17 | 6436 ± 139 <sup>a</sup> | 5864 ± 85 <sup>b</sup> | 6689 ± 158 <sup>a</sup> |
| PMC 1007.17 | 2949 ± 148 <sup>b</sup> | 1467 ± 88 <sup>c</sup> | 3655 ± 147 <sup>a</sup> |
| PMC 1008.17 | 1773 ± 95 | 1420 ± 103 | 1913 ± 192 |

**Table 2.** Recovery rate, limit of detection (LOD), and limit of quantification (LOQ) of the two tested extraction procedure (A and B, corresponding to pre-extraction or pre-analysis doping, respectively) measured on intestine, liver and muscle of the medaka fish. ^a^: determined on fresh weight; ^b^: determined on dry weight.

| Organ | Recovery rate |  | Limit of detection (LOD) |  | Limit of quantification (LOQ) |  |
| --- | --- | --- | --- | --- | --- | --- |
|  | Method A | Method B | µg.L<sup>-1</sup> | µg.g<sup>-1</sup> | µg.L<sup>-1</sup> | µg.g<sup>-1</sup> |
| Intestin<sup>a</sup> | 78 ± 13 % | 126 ± 22% | 0.412 | 20.6 | 1.25 | 62.5 |
| Liver<sup>a</sup> | 65 ± 7 % | 86 ± 15% | 0.506 | 2.53 | 1.25 | 62.5 |
| Muscle<sup>b</sup> | 25 ± 4 % | 51 ± 3% | 0.546 | 14.52 | 1.25 | 285 |

### Anatoxin-a toxicology

All individuals, including negative controls, gavaged with (±)-anatoxin-a doses up to 6.67 μg.g^−1^ survive without presenting any apparent symptoms of toxicosis and were able to recover from the tricaine sedation within less than 3 minutes when placed in fresh water. On contrary, all individual gavaved with 20 μg.g^−1^ (±)-anatoxin-a rapidly present (in the first 5 minutes) obvious signs of neurotoxic effects, comprising a complete stop or a rapid diminution of opercular movement, abnormal swimming with hemi- or complete paresis of the fins, accompanied with a global musculature rigidity. After 10 min, only one individual even presents few sporadic breathing activity, that completely stop after 15 min, when all other organisms already present complete paresis and ventilation cease. After 30 min, these individuals, presenting no sign of recovery, were considered as dying, if not dead, and euthanized, then were considered for further toxicological dose calculation. When LD_100_ and the NOAEL were observed at 20 μg.g^−1^ and 6.67 μg.g^−1^ (±)-anatoxin-a, respectively, the LD 50 was calculated as 11.5 μg.g^−1^ of (±)-anatoxin-a (Fig. 2A; table 3).

**Table 3.** Summary of lethal concentration estimators determined for anatoxin-a by different administration pathways on different vertebrate models.

| Model organism | Toxicological parameter | Toxicological values (µg.g<sup>-1</sup>) | Formulation of the toxicant | Reference |
| --- | --- | --- | --- | --- |
| Medaka fish | LD<sub>50</sub> by gavage | 11.5 | (±)-anatoxin-a | This study |
| Medaka fish | LD<sub>50</sub> by gavage | 5.75* | (+)-anatoxin-a | This study |
| Medaka fish | LD<sub>100</sub> by gavage | 20 | (±)-anatoxin-a | This study |
| Medaka fish | LD<sub>100</sub> by gavage | 10* | (+)-anatoxin-a | This study |
| Medaka fish | NOAEL by gavage | 6.67 | (±)-anatoxin-a | This study |
| Medaka fish | NOAEL by gavage | 3.33* | (+)-anatoxin-a | This study |
| Zebrafish | LD<sub>100</sub> by *i.p.* | 0.8 | (±)-anatoxin-a | Carneiro et al., 2015 |
| Mouse | LD<sub>50</sub> by *i.v.* | 0.1 | (+)-anatoxin-a | Fawell et al., 1999 |
| Mouse | LD<sub>50</sub> by gavage | [1-10] | (+)-anatoxin-a | Fawell et al., 1999 |
| Mouse | NOAEL by gavage | 3 | (+)-anatoxin-a | Fawell et al., 1999 |
| Mouse | LD<sub>50</sub> by gavage | 6.7 | *A. flos aquae* NRC 44-1 | Stevens et al., 1991 |
| Mouse | LD<sub>50</sub> by gavage | 16.2 | (+)-anatoxin-a | Stevens et al., 1991 |

\* determined according to the theoretical 1/1 (±)-racemic ratio, experimentally confirmed by LC-MS, considering as negligible the toxicity of (-)-anatoxin-a comparing to the 150-times higher toxicity of (+)-anatoxin-a (Osswald et al., 2007).

**Figure 2.**
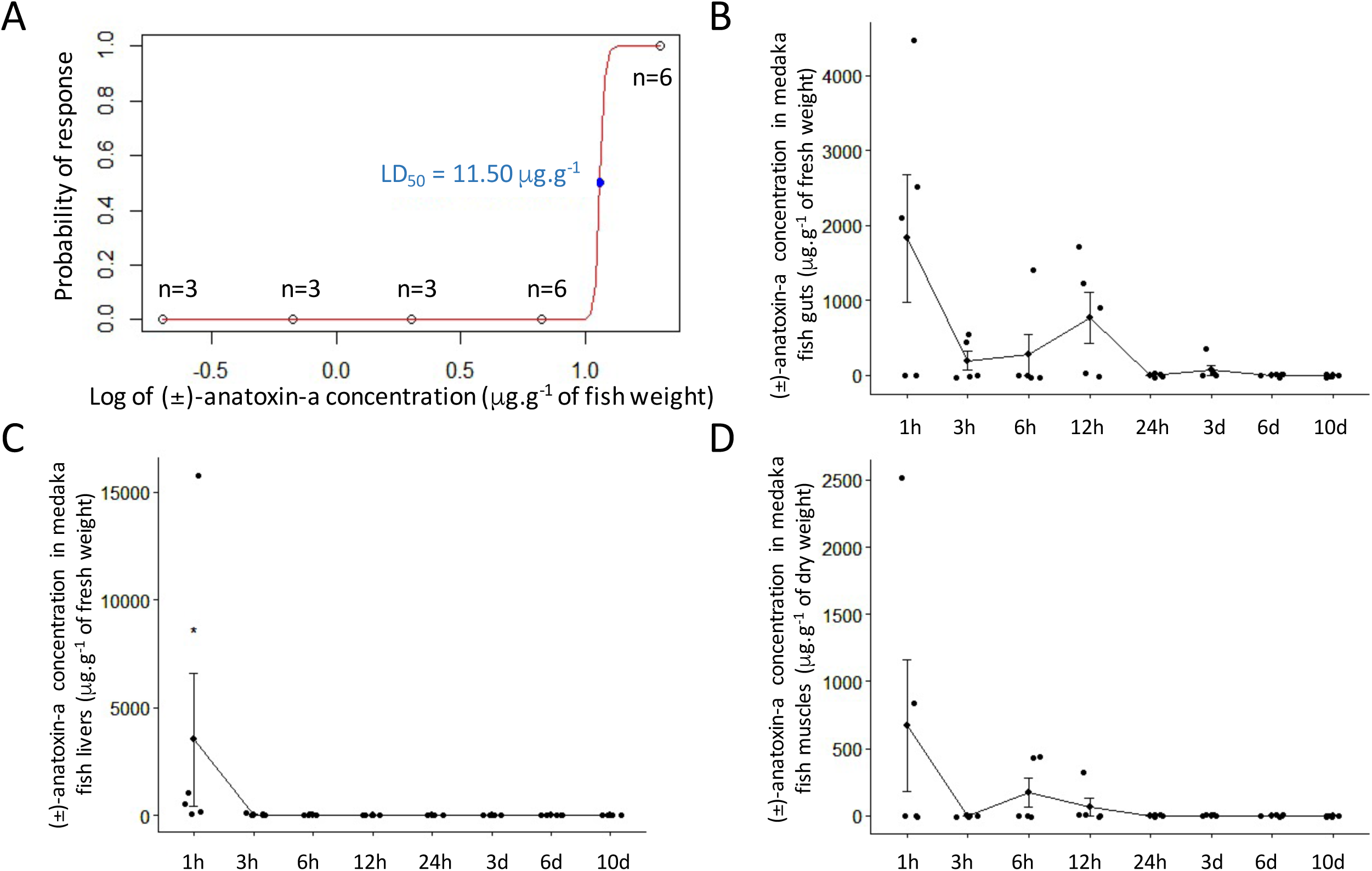
Anatoxin-a toxicity and toxico-kinetics on adult female medaka fish when administrated by single-dose gavage. Estimation of the LD_50_ of (±)-anatoxin-a by linear regression after log transformation (μg of (±)-anatoxin-a.g^−1^ of fish mass) (A). Toxico-kinetics of (±)-anatoxin-a administrated in a single NOAEL dose (6.67 μg.g^−1^ fish weight) in the gut (B), the liver (C) and the muscle (D) after 1h, 3h, 6h, 12h, 24h, 3j, 6j or 10j of depuration (n-5). On the 30 investigated individuals, only 6 of them does not present detectable amount of anatoxin-a, overall testifying for the global efficiency of the gavage experiments.

### Anatoxin-a toxico-kinetics

Following gavage experiment of adult female medaka fish to NOAEL, we have quantified (±)-anatoxin-a in the guts, the livers and the muscles after 1h, 3h, 6h, 12h, 24h, 3d, 6d and 10d, in order to monitor the dynamic of the anatoxin-a assimilation/depuration efficiency in these various compartments (Fig. 1B-D). The highest (±)-anatoxin-a amount was observed just 1h after the exposure, representing up to 15,789 μg.g^−1^ in some individual livers, representing more than 100 times more anatoxin-a than in the livers of other organisms similarly gavaged, illustrating the relative individual variability of the anatoxin-a uptake and assimilation/elimination during our experimentation. Although, fish tissues present a global individual variability, the larger amount of anatoxin-a were observed in livers, the guts, and in a lesser amount in the muscles after 1h, then the tissues present a rapidly decrease of anatoxin-a contents. Almost no more anatoxin-a was detectable after 24h in all tissues. The depuration rate was calculated for the 12 first hours of depuration as being of 57, 100 and 90%, in guts, livers and muscles, respectively, leading to a rapid elimination of the anatoxin-a that does no seems to bio-accumulate in any of the investigated fish tissues.

### Anatoxin-a effects on the liver metabolome

In order to investigate the molecular effects induced by (±)-anatoxin-a NOAEL exposure, the liver metabolite composition was compared between ungavaged fish (control) and fish collected 1h, 3h, 6h, 12h or 24h after gavage. The same livers extracts than those extracted with 75% methanol and analysed for anatoxin-a quantification were investigated by LC-MS/MS for untargeted metabolomics. A total of 591 different analytes were then extracted by the optimized pipeline, and their respective quantification (determined from the area-under-the-peak signal) compared between the different time-course groups using multivariate statistical methods, including unsupervised principal component analysis (PCA) and supervised partial least-squares discriminate analysis (PLS-DA), together with univariate groups variance analyses (ANOVA), in order to evaluate the potential of the method to discriminate among the experimental groups according to the time-course of anatoxin-a exposure and elimination.

Although the analysed fish livers present a global metabolome variability (Fig. 3A), the (±)-anatoxin-a gavage seems to rapidly modify the specific amount of various metabolites which progressively retrieved their initial state, as observed on components 1-3 projection of the PCA (Fig. 3B). The 29 analytes that present significant variation between the different groups according to ANOVA (P < 0.05) indicate a clear difference between the groups, as observed on heatmap with hierarchical clustering (Fig. 3D), with a rapid increase or decrease of the metabolite quantity that diminishes after few hour and almost recover control levels after 24h post-gavage (Fig. 3E). The supervised multivariate analysis (PLS-DA) model shows consistent R^2^ *cumulative*, Q^2^ *cumulative* and permutation scores (Fig. 3F). The most discriminating *m/z* features in the PLS-DA model (Figure 3C) were selected based on their respective VIP score, which resulted in 25 compounds with VIP value higher than 2 (Table 4), on component 1 and/or 2 (both contributing to the experimental group discrimination). The molecular formulas of each VIP were proposed based on accurate mass measurement, true isotopic pattern, and their putative identification were attempted with Metfrag and GNPS according to additional respective MS/MS fragmentation patterns. Interestingly, the tricaine (MW 165.0794 Da) belong to this VIP list and presents, as one could expect, a clear increase between the Control and 1h post-gavage fish livers (in relation with pre-gavage anesthesia procedure by balneation in 0.1% tricaine), then a complete disappearance between 1 and 3h.

**Table 4.**
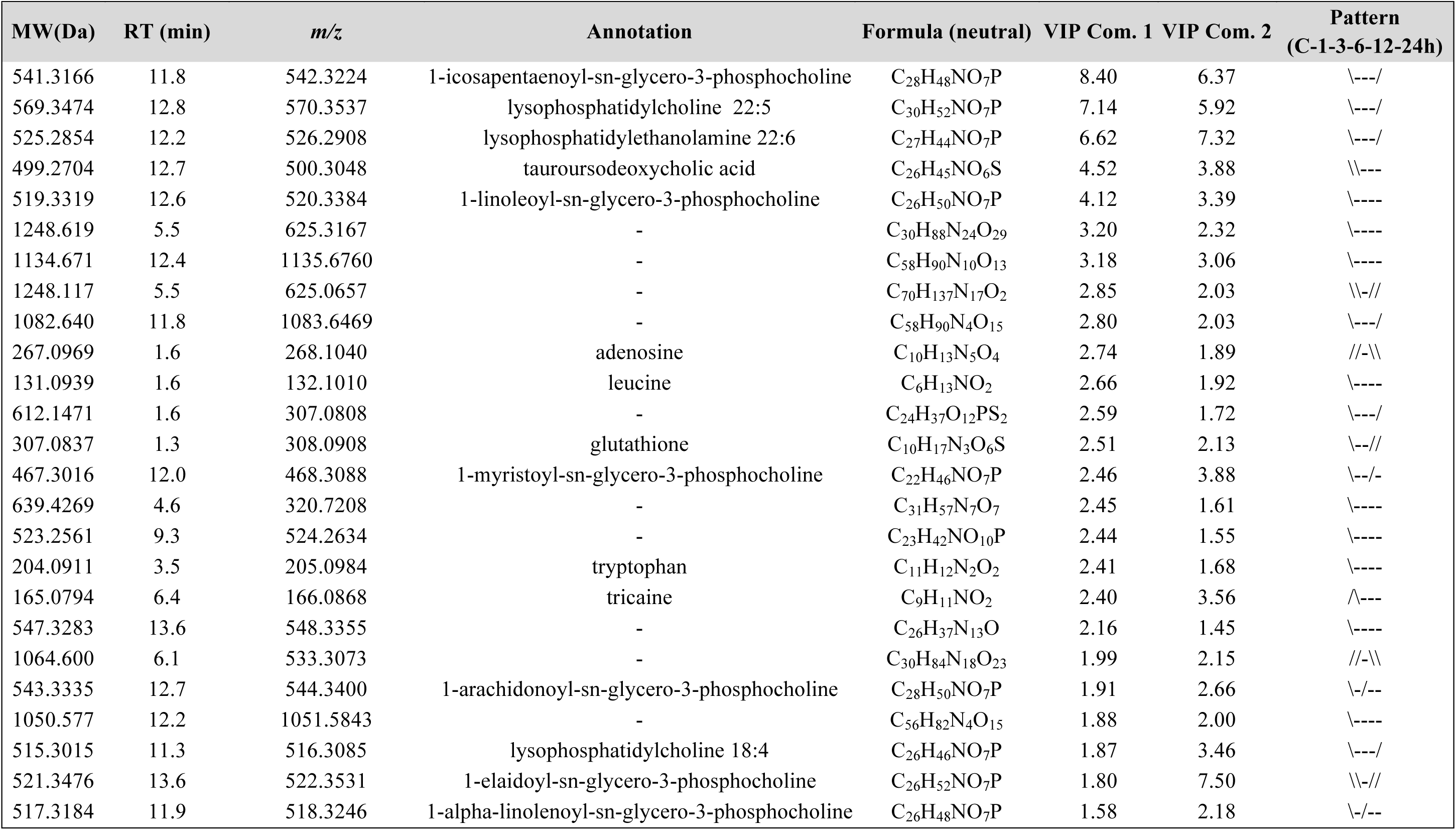
List of the 25 VIP analytes that present score > 2 on either component 1 or component 2 of the PLS-DA analysis performed with all treatment groups, and their putative annotation according to their respective high resolution mass, isotopic and MS/MS fragmentation patterns, searched against ChEBI, PubChem, HMDB and GNPS databases. “\”, “/” and “-” indicate when the metabolites globally decreases, increases or maintains their relative quantity between the time series, respectively.

**Figure 3.**
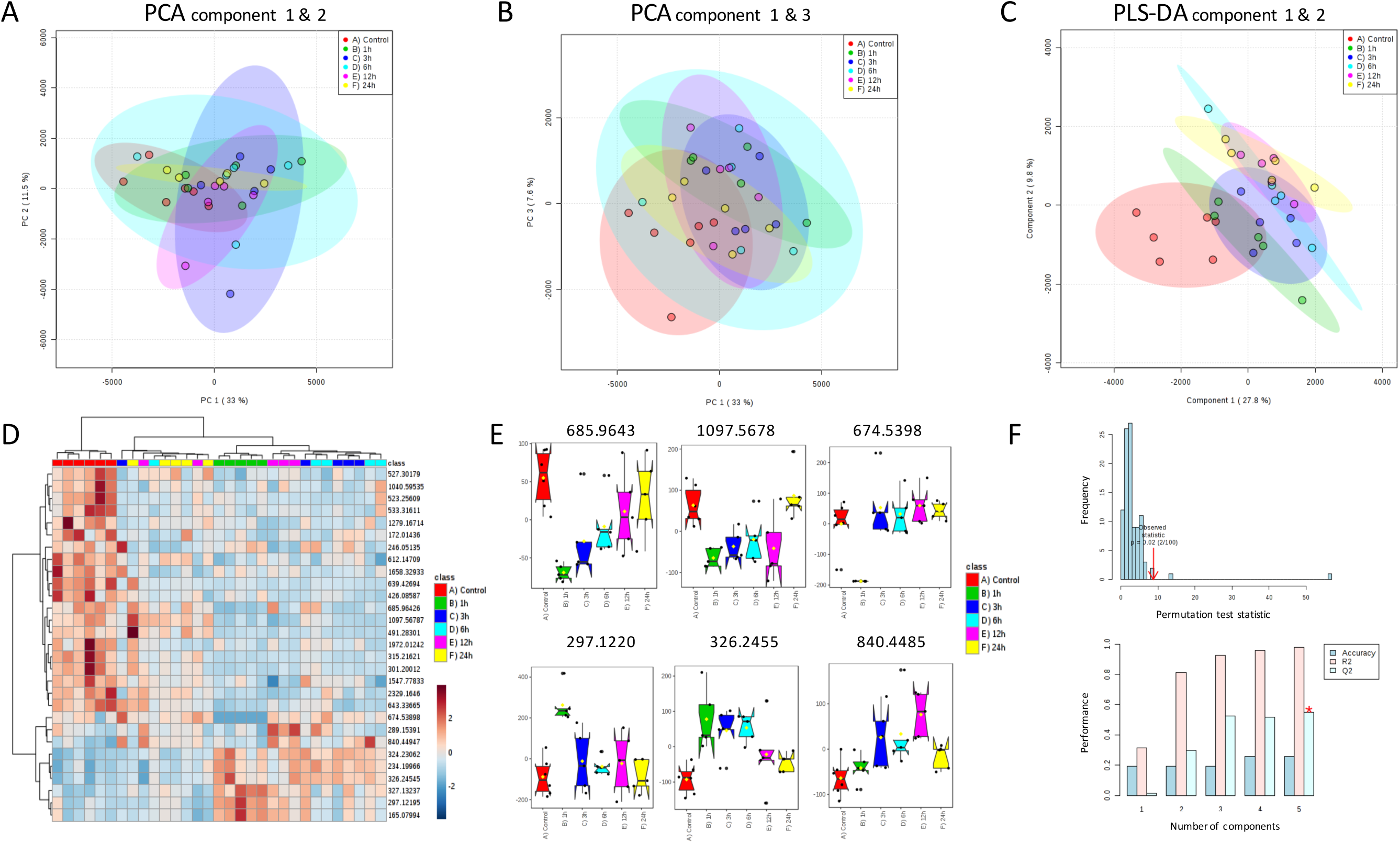
Metabolomics investigation of the liver metabolite variations after the administrated of a single NOAEL dose of (±)-anatoxin-a (6.67 μg.g^−1^ fish weight) by gavage of adult female medaka fish in the liver after 1h, 3h, 6h, 12h and 24h. Individual score plots generated from a principal component analysis performed with the intensity count of 591 variables according to components 1-2 (A) and 1-3 (B). Heatmap representation with hierarchical classification (Ward clustering according to Euclidian distances) performed from relative intensities of the 29 significantly dysregulated analytes (P <0.05 ANOVA) (D) and 6 examples of representative box-plots (E). Individual score plot generated from PLS-DA analysis performed with 591 extracted variables according to components 1-2 (C), and quality descriptors of the statistical significance and predictive ability of the discriminant model according to corresponding permutation and cross-validation tests (F), respectively.

The list of other putatively annotated compounds comprises then various phospholipids (n=9) belonging to the glycerol-phosphocholine group, all presenting comparable variation patterns among the experimental groups, with an initial drop of these metabolite quantity between the control and the 1h group, then a progressive re-increase until an almost complete recovery of the initial metabolite amount within less than 24h. Those metabolites are directly related to lipid metabolism process and their successive decrease and increase within the liver may denote an important lipid consumption by the organism and a progressive recover of the liver to an unstressed condition. On the contrary, the sole VIPs which relative quantities present an initial increase were an undetermined analyte (MW 1064.600 Da) and the adenosine, both retrieving their initial levels within less than 24h. Such adenosine transient increase may denote an intensification of hepatic blood circulation, as it presents direct effect on vasodilation of liver arteria (Robson & Schuppan 2010). Although our experimental design does allow to discriminate the specific effects of the anatoxin-a from those of the tricaine anesthesia alone, it overall shows that the organisms present a complete recovery 24h after being gavaged with anatoxin-a single NOAEL dose.

## Discussion

In the present study, the neurotoxicological response of medaka fish subjected to anatoxin-a appears comparable to that found in carp (Osswald et al., 2007a) and zebrafish (Carneiro et al., 2015) supporting the evidences for the existence of a similar mechanism of action. Whereas no apparent precursor effect appears when fish are gavaged with sub-acute doses of anatoxin-a, the symptoms observed in the fish exposed to a higher dose denote an all-or-nothing effect, comprising rapid and intense neurotoxic signs, that are compatible with the mechanism of toxic action of anatoxin-a in the nervous system that has been described so far for other vertebrates (Fawell et al., 1999). These symptoms comprise different manifestations of muscular paralysis that are likely due to the primary, and nearly irreversible, binding of the anatoxin-a, being an acetylcholine antagonist, to the acetylcholine receptors of the cholinergic synapses of the neuromuscular junctions, leading to a continuous muscular contraction. Previous works indicate that below a certain levels the effects appear to be transient, the animals being able to make a complete and rapid recovery, although the data available in the literature remains limited (Dittman and Wiegand 2006).

Our dose-response toxicological analysis indicates that anatoxin-a exhibits toxicological reference dose of 5.75 and 3.33 μg.g^−1^ of medaka body weight for LD_50_ and NOAEL, respectively. This results are in remarkable agreement with precedent data obtained on other organisms. Indeed, previous investigations of anatoxin-a toxicity determined on mouse that, when administrated by single dose gavage, (+)-anatoxin-a exhibits a LD_50_ and a NOAEL of above 5 and 3 μg.g^−1^ of body weight, respectively (Fawell et al., 1999). In addition, Stevens and co-workers (1991) have initially shown that (+)-anatoxin-a presents LD_50_ value of 16.2 and 6.7 μg.g^−1^ of body weight, for pure (+)-anatoxin-a and complex extract containing anatoxin-a administrated by single gavage to mouse. Taken together, those results suggest that medaka fish and mouse apparently present very similar toxicological dose-response to anatoxin-a. Indeed, as the unnatural (-)-anatoxin-a isomer was shown to insignificantly contribute to the global toxicity to the 1/1 racemic mixture of (±)-anatoxin-a used in this study (MacPhail et al., 2007), one could estimate that toxicity values of (+)-anatoxin-a might theoretically be very similar for medaka fish than for mouse.

In our experiments, the promptitude of the neuromuscular effects of anatoxin-a when administrated by gavage are in agreement with previous investigation performed on mice (Fawell et al., 1999), suggesting that anatoxin-a might be rapidly assimilated by the organisms by crossing the intestine barrier and being widespread to the whole musculature within less than 2 minutes. When anatoxin-a producing cyanobacterial biofilm are accidently ingested by dogs, the animals present first appearance of neurotoxic symptoms within less than 5 minutes (Wood et al., 2007), testifying for a very rapid assimilation of a toxinogenous dose of anatoxin-a released from the cyanobacterial biomass within the stomach. To date, the genuine mechanism of the anatoxin-a transfer through the intestinal epithelia remains undetermined but the promptitude of the effects suggests that this small molecule (MW = 165 da) may remain uncharged in order to be able to sharply cross the intestinal barrier, potentially through passive paracellular diffusion (Dahlgren and Lennernäs 2019).

Our toxico-kinetics investigation has shown that no detectable amount of anatoxin-a was still observed in guts 12h after the fish having been gavaged to a single NOAEL dose. Then, the medaka fish presents also a rapid elimination of the anatoxin-a that transiently have been addressed to the liver, and in a lesser extent to the muscle, indicating that it may not being apparently accumulating anatoxin-a under those conditions. Interestingly, Osswald and co-workers (2011) have shown no significant bioaccumulation of anatoxin-a in the trout tissues when administrated by balneation up to 5 mg.L^−1^ of anatoxin-a. Previously, Osswald et al. (2008) have also similarly shown no bioaccumulation of anatoxin-a by the Mediterranean mussel *M. galloprovincialis*, that has been otherwise shown to present high accumulation capability for various contaminants, such as microcystins (Vasconcelos 1995). Although they observed some anatoxin-a uptake from the surrounding water filtered by the animals (observed maximum accumulation efficiency = 11%), this seemed rapidly reduced rather by the depuration process of phase II detoxification enzymes or by passive elimination processes. These data show that, as anatoxin-a is capable to rapidly penetrate the whole organisms body, and induces obvious and acute neurotoxicological effects within a minute, it may also be rapidly eliminated by classical excretion or depuration mechanisms.

## Conclusion

In our gavage experiment of medaka fish to a single NOAEL dose of anatoxin-a, the toxin appears to have been rapidly eliminated and the molecular effects were no more perceptible within the fish liver metabolome after 24h.

These observations suggest that when the dose remains below the acute toxicological limit producing neurotoxicosis that can lethal consequences, the organism are able to make a completely and rapidly recovery, that seems not to induce obvious effects even if the exposure is repeated several times (Fawell et al. 1999). However, one could still suspect that chronic anatoxin-a exposure may induce more insidious long-term pathologies on the central nervous system (Lombardo and Maskos 2015). But this hypothesis still remains to be explored.

More over, the accurate investigations of the fish flesh contamination by anatoxin-a under highest toxinogenous cyanobacterial proliferations, especially when, in nature, certain fish are potentially actively feeding on those biofilms (Ledreux et al., 2014), still remain to be performed in order to provide convincing data supporting the evaluation of the associated risks.

## Acknowledgements

This work was supported by grant CRD from ANSES attributed to the Cyanariv project, lead by Catherine Quiblier.

*The authors declare not conflict of interest*

